# The collective influence of 1, 25-dihydroxyvitamin D_3_ with physiological fluid shear stress on osteoblasts

**DOI:** 10.1101/146159

**Authors:** Yan Li, Jiafeng Yuan, Qianwen Wang, Lijie Sun, Yunying Sha, Yanxiang Li, Lizhong Wang, Zhonghua Wang

## Abstract

1, 25-dihydroxyvitamin D_3_ (1, 25 (OH)_2_ D_3_) and mechanical stimuli in physiological environment play an important role in the pathogenesis of osteoporosis. The effects of 1, 25-dihydroxyvitamin D_3_ alone and mechanical stimuli alone on osteoblasts have been widely investigated. This study reports the collective influences of 1, 25-dihydroxyvitamin D_3_ and flow shear stress (FSS) on biological functions of osteoblasts. 1, 25 (OH)_2_ D_3_ were constructed in various kinds of concentration (0, 1, 10, 100 nmmol/L), while physiological fluid shear stress (12 dynes/cm^2^) were produced by using a parallel-plate fluid flow system. 1, 25 (OH)_2_ D_3_ affects the responses of ROBs to FSS, including the inhibition of NO releases and cell proliferation as well as the promotion of PGE_2_ releases and cell differentiation. These findings provide a possible mechanism by which 1, 25(OH)_2_ D_3_ influences osteoblasts responses to FSS and may provide guidance for the selection of 1, 25(OH)_2_ D_3_ concentration and mechanical loading in order to *in vitro* produce functional bone tissues.

## 1. Introduction

1, 25-dihydroxyvitamin D_3_ (1, 25 (OH)_2_ D_3_) is a kind of vitamin D active metabolite, which plays an important role in the pathogenesis of osteoporosis. The features of osteoporosis are low bone mass and bone microstructure destruction. Osteoporosis is a systemic metabolic bone disease and increased bone fragility and susceptibility to fracture. For the treatment of osteoporosis, vitamin D active metabolite has been applied widely. What’s more, 1, 25-dihydroxyvitamin D_3_ is one of the most widely applied vitamin D active metabolite and plays an significant role in the regulation of calcium and phosphate metabolism, also can obviously affect the proliferation, differentiation of osteoblasts^[1–3]^. The researchers found that 1, 25 (OH)_2_ D_3_ inhibits cell growth and enhances osteoblast differentiation by regulating the expression of collagenous and non-collagenous proteins, including osteocalcin and osteopontin^[4, 5]^. Further, M. Van Driel and V.J.Woeckel demonstrated that 1, 25 (OH)_2_ D_3_ can enhance the mineralization of osteoid tissue produced by osteoblasts^[6, 7]^.Thus, Understanding the effects of 1, 25 (OH)_2_ D_3_ on the perception of osteoblasts is beneficial for the treatment of osteoporosis, inducing appropriate cell responses and promoting bone formation.

In addition, bone is a kind of porous dynamic tissue in the body. It is always subjected to a variety of mechanical stimulation from outside and inside. If the receiving force of bone is reduced, such as bedridden for a long time, bone fixation or a decrease in physical activity, which leads to increase the absorption of bone, severe bone loss, and enhance the risk of osteoporosis. The structure of bone may be caused and maintained by the difference between the actual receiving mechanical stimulation of bone and the mechanical amplitude of normal internal environment. At the level of cells, fluid shear stress (FSS) generated in canalicular systems of physiological bone is regarded as the principal mechanical stimulus responsible for bone adaption and remodeling^[8–10]^. As a result, FSS is widely employed as an external mechanical stimulus in bioreactors for bone tissue engineering^[1, 12]^. Various magnitudes of FSS from 10 dynes/cm^2^ to 20 dynes/cm^2^ have been concluded to influence the morphological responses, proliferation and differentiation of osteoblasts with a magnitude-dependent and duration-dependent manner^[13–16]^. Particularly, Liu et al^[16]^ reported that stimulation of 16 dynes/cm^2^ or 19 dynes/cm^2^ FSS for 1h induced osteoblasts reorientation along FSS direction while 12 dynes/cm^2^ FSS did not. Kapur^[13]^ found that 20 dynes/cm^2^ FSS for 30min obviously promoted osteoblasts proliferation and differentiation, while Horikawa^[17]^ pointed out that long-term FSS stimulation such as 10 dynes/cm^2^ for over 1h had destructive effects on osteoblasts. In bone tissue *in vivo*, the physiological FSS caused by interstitial fluid flow is 8~30 dynes/cm^2^ on the basis of the model presented by Weinbaum^[18]^. In this study, 12 dynes/cm^2^ were employed to represent physiological FSS.

The aim of this study is to investigate the collective effects of 1, 25 (OH)_2_ D_3_ with physiological fluid shear stress on osteoblasts. To this end, 1, 25 (OH)_2_ D_3_ were constructed in various kinds of concentration (0, 1, 10, 100 nmol/L), while physiological fluid shear stress (12 dynes/cm^2^) were produced by using a parallel-plate fluid flow system. The plasma membrane integrity, the short-term response on releases of nitric oxide (NO) and prostaglandin E_2_ (PGE_2_) and the long-term response on cell proliferation (Proliferation index, PI), alkaline phosphatase (ALP), osteopontin (OPN) and osteocalcin (OCN), these all were selected to characterize the responses of osteobalst to collective stimulation. To elaborate the potential mechanism, the focal adhesions formation (FA) and cytoskeleton rearrangement (F-actin) before and after FSS stimulation were further examined. The cell morphology and orientation were quantified as well. Selection of NO and PGE_2_ releases is based on the fact that they are the early rapid responses of osteoblasts to FSS and play important roles in bone formation and remodeling^[19–22]^.

## 2. Materials and methods

### 2.1. Osteoblasts culture

Primary rat osteoblasts (ROBs) cultures were described in our previous paper^[23]^. Briefly, the calvaria bone of SD rats was removed and placed in petri dish containing PBS buffer. The periosteum and surrounding connective tissue were removed by using tweezers. The calvaria bone was washed with PBS and cleaned with DMEM until the surface of bone is white and transparent, immersed into a small amount of fetal calf serum. The calvaria bone was cut into about 1mm×1mm×1mm pieces, coated uniformly in 25ml culture flask. Cultures were initiated in DMEM (Gibico, USA) supplemented with 10% heat-inactivated fetal calf serum (FCS, Sijiqing, China), penicillin (100U/mL), streptomycin (100μg/mL), and 0.05% L-glutamine, and maintained in a humidified atmosphere of 5% CO_2_/95% air at 37°C. The medium was changed every two days. After confluence, the cells were sub-cultured and identified by using the von Kossa staining method according to previously reported procedure^[23]^. The fourth to sixth passage of ROBs were used for experiments at a density of 2×10^5^ cells/slide.

### 2.2. Physiological fluid shear stress and static treatments

A parallel-plate flow chamber apparatus was employed to provide FSS^[24]^. The dimension of the flow chamber was 7.50cm (length, L) × 2.50 cm (width, W) × 0.03cm (height, H). FSS was produced by circulating 10 mL DMEM using a peristaltic pump (JieHeng, China). The produced FSS (τ, dynes/cm^2^) was calculated according to equation *τ = 6ηQ* / *H*^2^W, where *η* is the dynamic viscosity of the perfusate and *Q* is the flow rate. To study the collective effects of 1, 25 (OH)_2_ D_3_ and FSS on the responses of osteoblasts, pre-incubations of ROBs for 24h were performed in medium with 1, 25 (OH)_2_ D_3_ (0, 1, 10, 100 nmol/l) (Sigma-Aldrich, USA) under static conditions. The concentrations of 1, 25 (OH)_2_ D_3_ were chosen based on a previous study in which the different concentration of 1, 25 (OH)_2_ D_3_ affected osteoblast proliferation and differentiation to a similar extend^[5, 25, 26]^. Thereafter, osteoblasts were subjected to FSS or static treatment asdescribed previously^[27]^. Shortly, the employed physiological FSS in this study was 12 dynes/cm^2^. All the components of the apparatus, except the pump, were maintained in a 37°C incubator during the experiment and the medium was continuously saturated with 5% CO_2_/95% air. For FSS-loaded samples (denoted by X-FSS, where X represents different concentration of 1, 25 (OH)_2_ D_3_), slides with attached ROBs were first mounted on the flow chamber and then exposed to FSS for a predetermined time. Other samples without FSS exposure (denoted by X-Static) were kept in Petri dishes at 37°C in a humidified atmosphere of 5% CO_2_/95% air for the same predetermined time.

### 2.3. Determination of NO, and PGE_2_ releases

After incubation with 1, 25 (OH)_2_ D_3_ for 24h, ROBs were employed for the determination of NO, and PGE_2_ releases. After ROBs on slides were exposed to FSS for a predetermined time, 2 mL of medium was withdrawn and replenished with an equal volume of fresh culture medium to maintain a constant circulating fluid volume. The same procedure was performed for samples without FSS exposure.

#### 2.3.1 NO release

NO concentration was detected by using a Nitric Oxide Assay Kit (Beyotime, China) based on Griess Reagent. The withdrawn medium was permitted to react with Griess Reagent for 10 min, and then the absorbance was measured at 540nm using a Microplate Reader (Bio-Rad, USA). The NO concentration was determined according to the standard curve plotted by the supplied NaNO_2_ standards and normalized to the total cellular protein.

#### 2.3.2 PGE_2_ release

PGE_2_ release was measured by using a Rat PGE_2_ ELISA Assay Kit (Yuan-Ye Chemical reagents, China). The absorbance was detected at 450nm and the PGE_2_ concentration was determined on the basis of the standard curve plotted by the supplied PGE_2_ standards and normalized to the total cellular protein.

### 2.4. Determination of the proliferative index (PI)

After being exposed to FSS for 1 h, the cells were trypsinized, washed with PBS, and fixed in 70% ethanol at 4°C. The cell suspension was centrifuged at 2000r/min for 10min. The collected ROBs were washed with PBS and then 1mL of PBS was added to obtain the cell suspension again. Following this, 20 μL of RNAse (10 mg/mL) was added to the obtained cell suspension and it was then incubated at 37°C for at least 30min. Then, 50μL of pyridine iodide was added to the cell suspension and it was incubated in the dark at 37°C for another 30 min. Finally, the cell cycle stages were detected using a Flow Cytometry (BD FACS Vantage SE, USA) and analyzed using Cell quest software (Becton Dickinson). The cell proliferative index (PI%) was calculated using the following formula: PI% = [(S + G2/M) /(G0/G1 + S + G2/M)]%, where S, G2/M, and G0/G1 represent the cell numbers in phase S, phase G2 and M, and phase G0 and G1, respectively, in cell cycle.

### 2.5. Plasma membrane integrity after FSS exposure

Lactate dehydrogenase (LDH) level released from cytoplasma is a common parameter to indicate the integrity of plasma membrane^[28]^. After exposure of ROBs to FSS for 1 h, 500 μL of the conditioned medium was employed for detection of LDH level by using a LDH Assay Kit (Jiancheng, China). The basic principle of this assay is that NADH, generated by oxidization of lactate into pyruvate under the catalysis of LDH, may convert iodonitrotetrazolium into a red formazan product in the presence of diaphorase. The absorbance of formazan at 490 nm is proportional to the amount of LDH in the medium. Comparison of the LDH level with FSS exposure to that without FSS exposure may indicate the integrity of plasma membrane.

### 2.6. Focal adhesions and cytoskeletons of ROBs before and after FSS exposure

Focal adhesion formation and cytoskeleton rearrangement of ROBs before and after 1h FSS exposure were examined by immunofluorescence staining. Before staining, ROBs were fixed through incubation in 4% (w/v) paraformaldehyde in PBS for 30 min and permeabilized for 10 min with 0.25% (v/v) Triton X-100 in PBS. Subsequently, 1% BSA in PBS was added to block the non-specific binding sites by incubating with ROBs for 1h. To observe focal adhesion formation, vinculin was stained by initially incubating with a mouse monoclonal anti-vinculin antibody at room temperature for 60 min, and subsequently incubating with a FITC-conjugated anti-mouse immunoglobulin secondary antibody at room temperature for another 60min. F-actin was stained using Rhodamine-Phalloidin at room temperature for 60 min. Finally, the nuclei were stained by incubation with bisBenzimide H 33258 at room temperature for 10min. The stained specimens were washed, mounted in glycerol, and examined by a confocal laser scanning microscope (CLSM; TCS SP5, Leica, Germany).

### 2.7. ALP assay

ROBs were exposed to FSS for 1 h, the ROBs were collected and lysed for 30 min in 150 μL lysis buffer (Cwbiotech, China) on ice. The total proteins were got in supernatants with centrufugation (14000 r/min). The ALP activity was detected by ALP assay kit (Nanjing Jiancheng Bioengineering Institute, China). The total protein concentration, detected by BCA Protein Assay Kit (Beyotime, China), was used to normalize ALP activity. The absorbance was detected by Multifunctional Microplate Reader (Bio-Rad) at 490 nm and 570 nm for ALP and BCA analysis, respectively.

### 2.8. Quantitative real-time polymerase chain reaction (qRT-PCR)

After exposure to FSS for 1 h, the differentiation of osteoblasts stimulated with both 1, 25 (OH)_2_ D_3_ and FSS was further assessed by qRT-PCR. The relative mRNA expression levels of commonly used bone markers, including osteopontin (OPN) and osteocalcin (OCN) (primer pairs used are shown in Table 1), were measured. Total RNA isolation for qRT-PCR samples was conducted using an RNA simple Total RNA Kit (Biotake, Beijing, China). Complementary DNA (cDNA) was then reversetranscribed from 1 mg of total RNA using a RevertAid First Strand cDNA Synthesis Kit (ThermoFisher Scientific, USA) according to manufacturer instructions. Realtime PCR was performed on SYBR Green PCR Master Mix (T advanced, JENA, Germany) and the reaction was carried out using an ABI 7500 system (Scandrop, JENA, Germany). Finally, housekeeping gene glyceraldehyde-3-phosphate dehydrogenase (GAPDH) was used to normalize therelative mRNA expression level of each gene and quantification was based on the cycle threshold (CT) values.

**Table 1.**
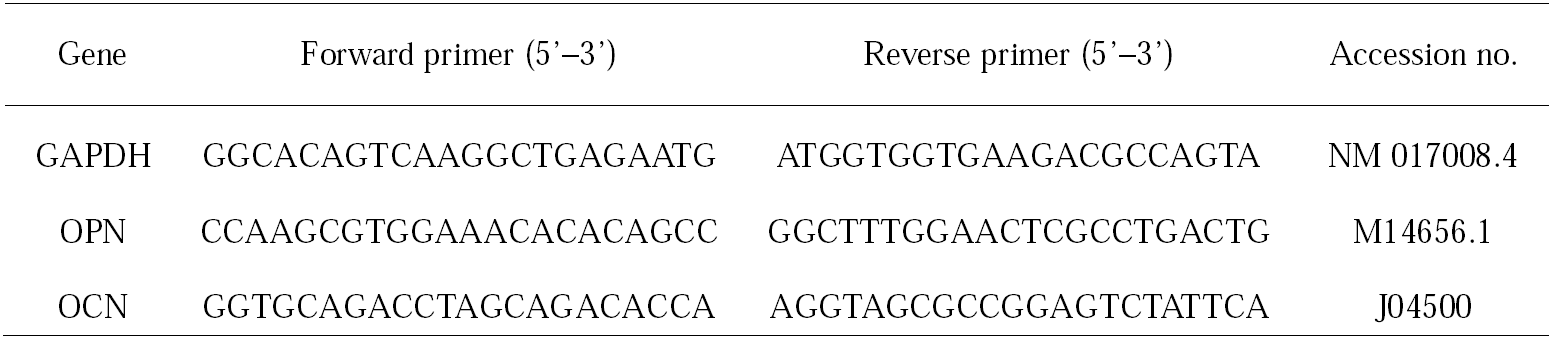
OPN and OCN used in PCR analysis

### 2.9. Statistical analysis

Data were expressed as means ± SD (n ≥6 for all experiments). Single factor analysis of variance (ANOVA) technique with OriginPro (version 8.0) was used to assess the statistical significance of results. The confidence interval was set to 0.05.

## 3. Results

### 3.1. NO and PGE_2_ releases

First, we examined the effects of 1, 25 (OH)_2_ D_3_ alone on the NO and PGE_2_ releases of ROBs. As shown in Fig. 1, when no FSS was applied, ROBs released a small amount of NO (Fig. 1) and PGE_2_ (Fig. 3). All the releases demonstrated no observable change with incubation time. This implies that 1, 25 (OH)_2_ D_3_ alone has negligible effects on NO and PGE_2_ releases of ROBs. When FSS was applied, the NO and PGE_2_ releases were significantly increased compared to their corresponding static counterparts (Fig. 1-3). Nevertheless, 10-FSS and 100-FSS had and obvious decrease in NO releases compared to the 0-FSS and 1-FSS, also 0-FSS ≈ 1-FSS (Fig. 2). But the opposite phenomena were observed for PGE_2_ releases (Fig. 3). It is seen that FSS induced significant increase PGE_2_ releases (Fig. 3). This suggests that FSS-induced NO and PGE_2_ releases of ROBs are 1, 25 (OH)_2_ D_3_ concentration dependent with a pattern of 0-FSS > 1-FSS > 10-FSS > 100-FSS for NO (Fig. 2), and 100-FSS > 10-FSS > 0-FSS ≈ 1-FSS for PGE_2_ (Fig. 3).

**Fig. 1.**
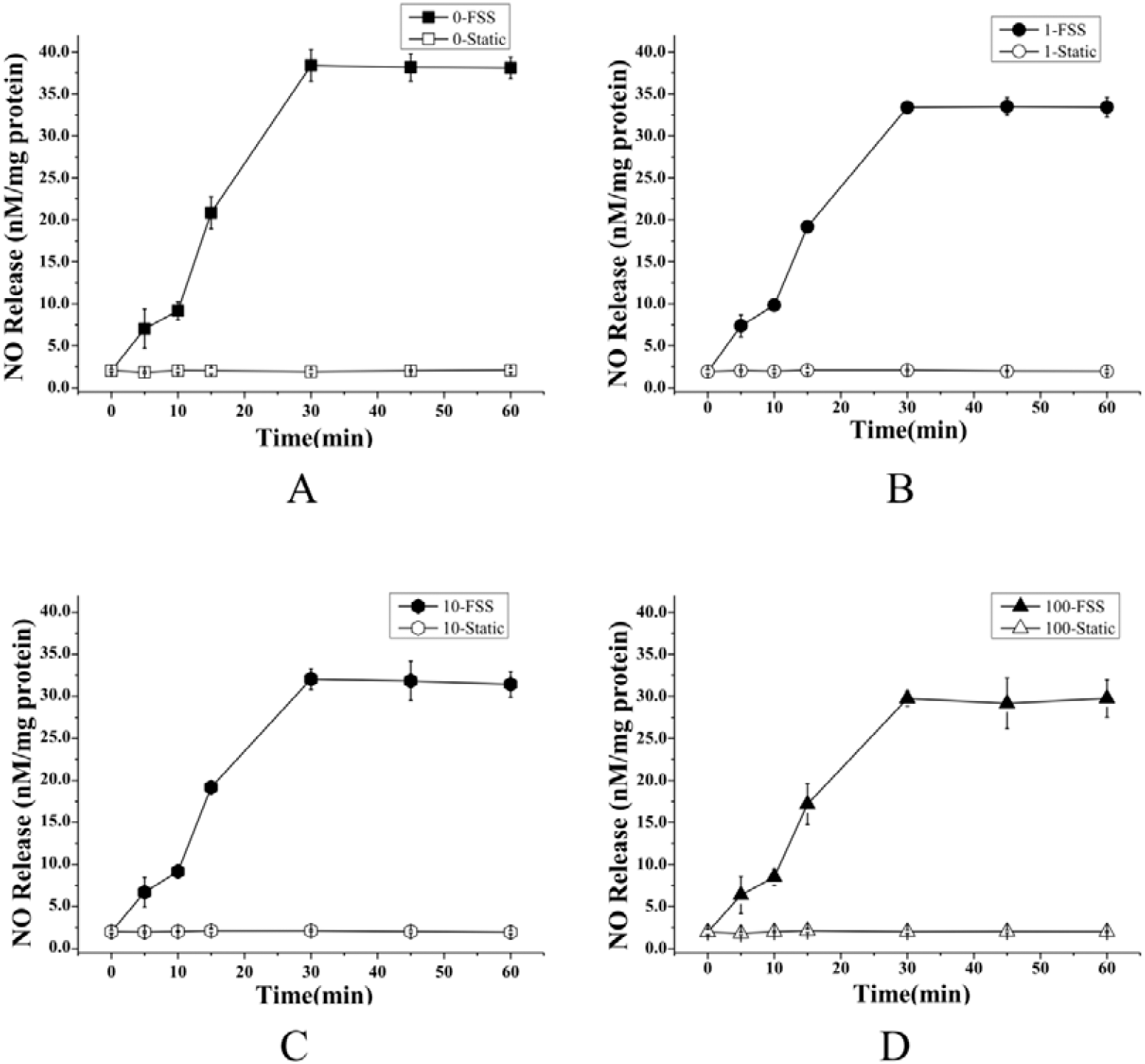
Releases of NO with different 1, 25 (OH)_2_ D_3_ concentration for Static and FSS.

**Fig. 2.**
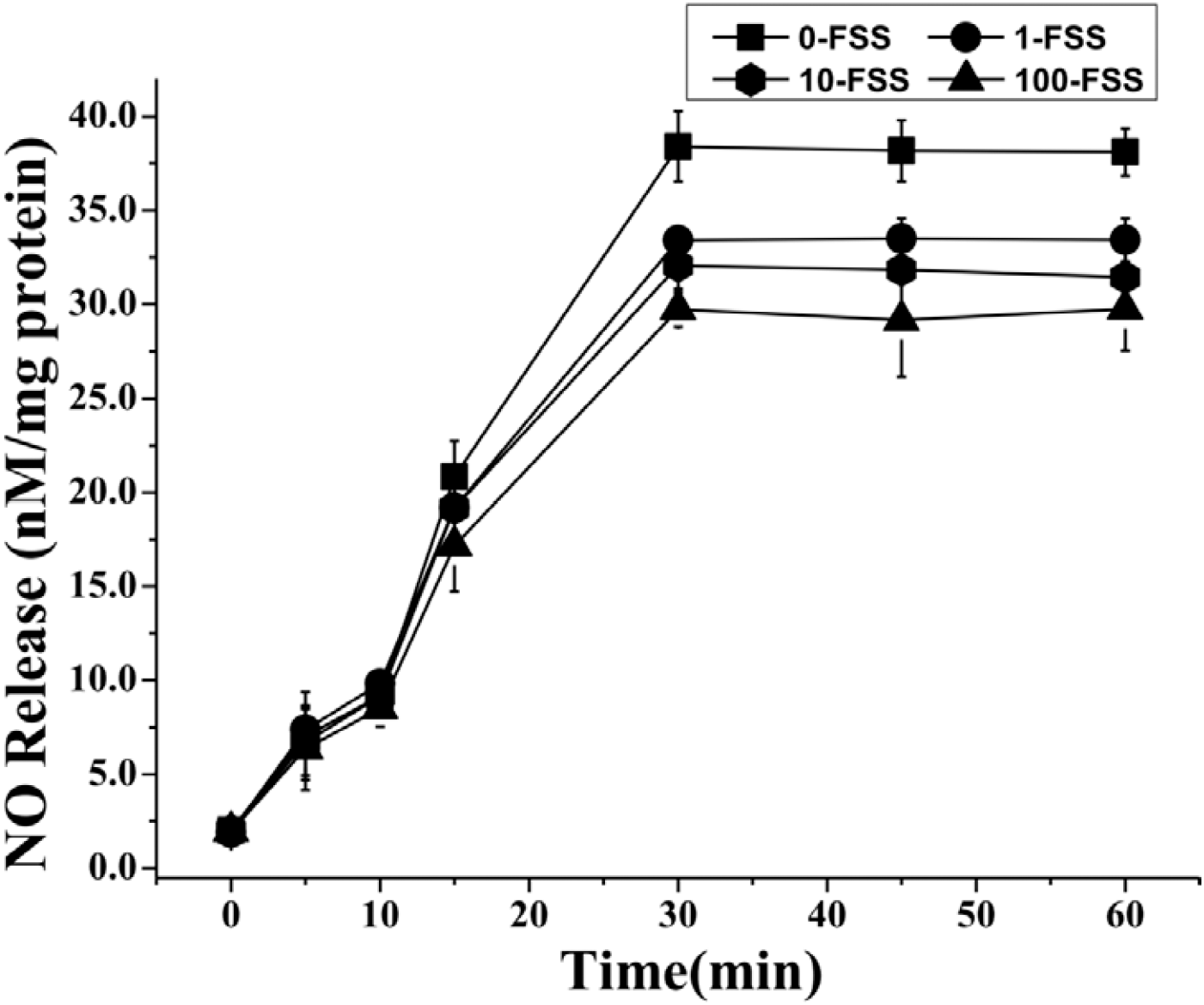
Releases of NO with different 1, 25 (OH)_2_ D_3_ concentration for FSS.

**Fig. 3.**
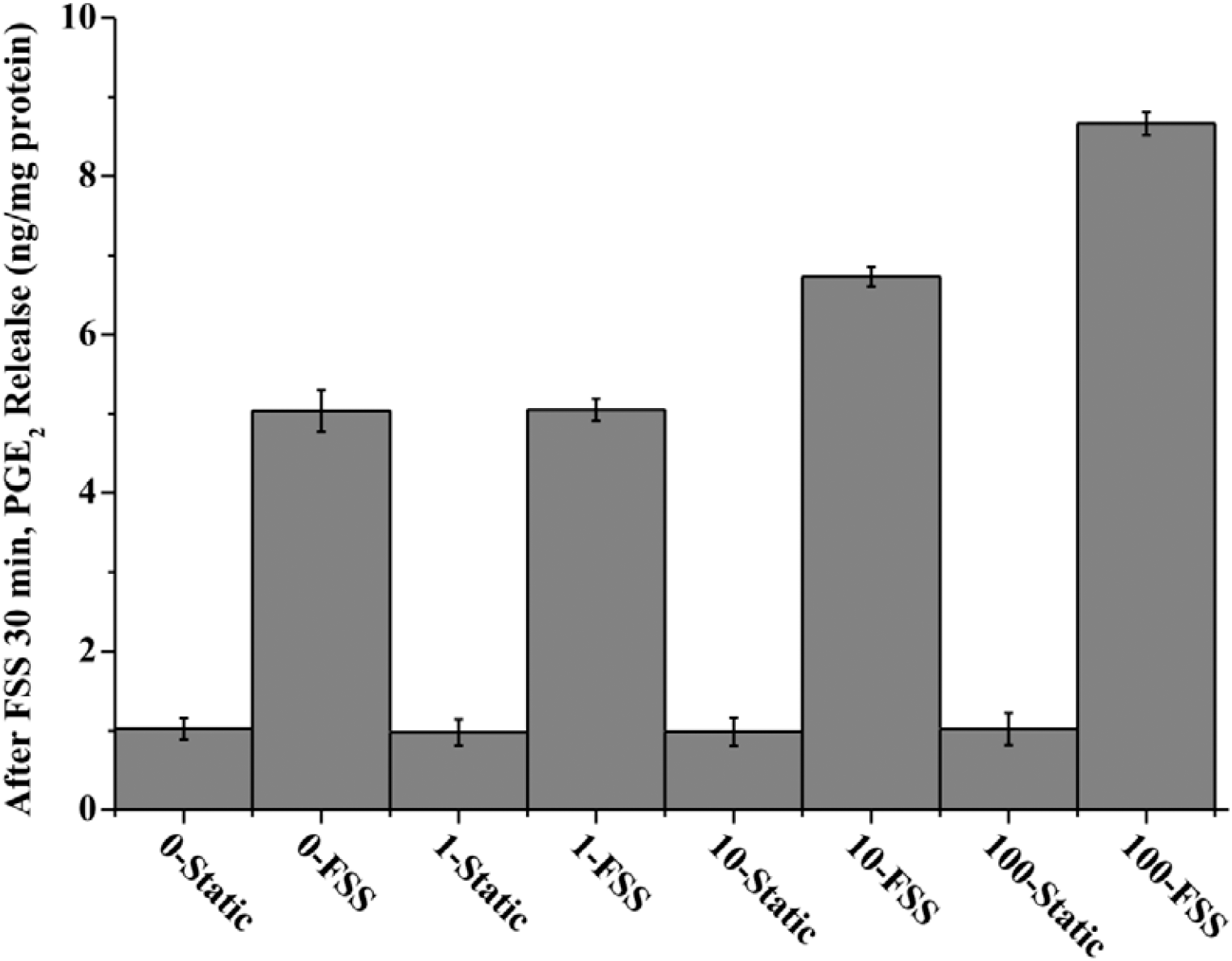
Releases of PGE_2_ with different 1, 25 (OH)_2_ D_3_ concentration for Static and FSS.

### 3.2. Determination of the proliferative index (PI)

The cell proliferation potential can be indicated by cell proliferative index (PI), which is the ratio of cell number in G2/M phase and S phase to the total cell number. The PI values of ROBs incubated by 1, 25 (OH)_2_ D_3_, without or with FSS exposure, were determined by using flow cytometry and the result is illustrated in Fig. 4. The typical histograms of the PI cell-cycle profiles are shown in Fig. 5. It is clear that 1, 25 (OH)_2_ D_3_ inhibited ROBs proliferation, and the stronger concentration of 1, 25 (OH)_2_ D_3_, the stronger inhibition. But after applying FSS, the PI of all X-FSS is increased, and higher than those of 0-Static, which indicates that the exposure of FSS counteracts the inhibitory effect of 1, 25 (OH)_2_ D_3_ on ROBs proliferation. The PI values for ROBs exposed to various concentrations of 1, 25 (OH)_2_ D_3_ and FSS were shown in Fig. 4 and followed the pattern 0-FSS > 10-FSS > 1-FSS > 100-FSS.

**Fig. 4.**
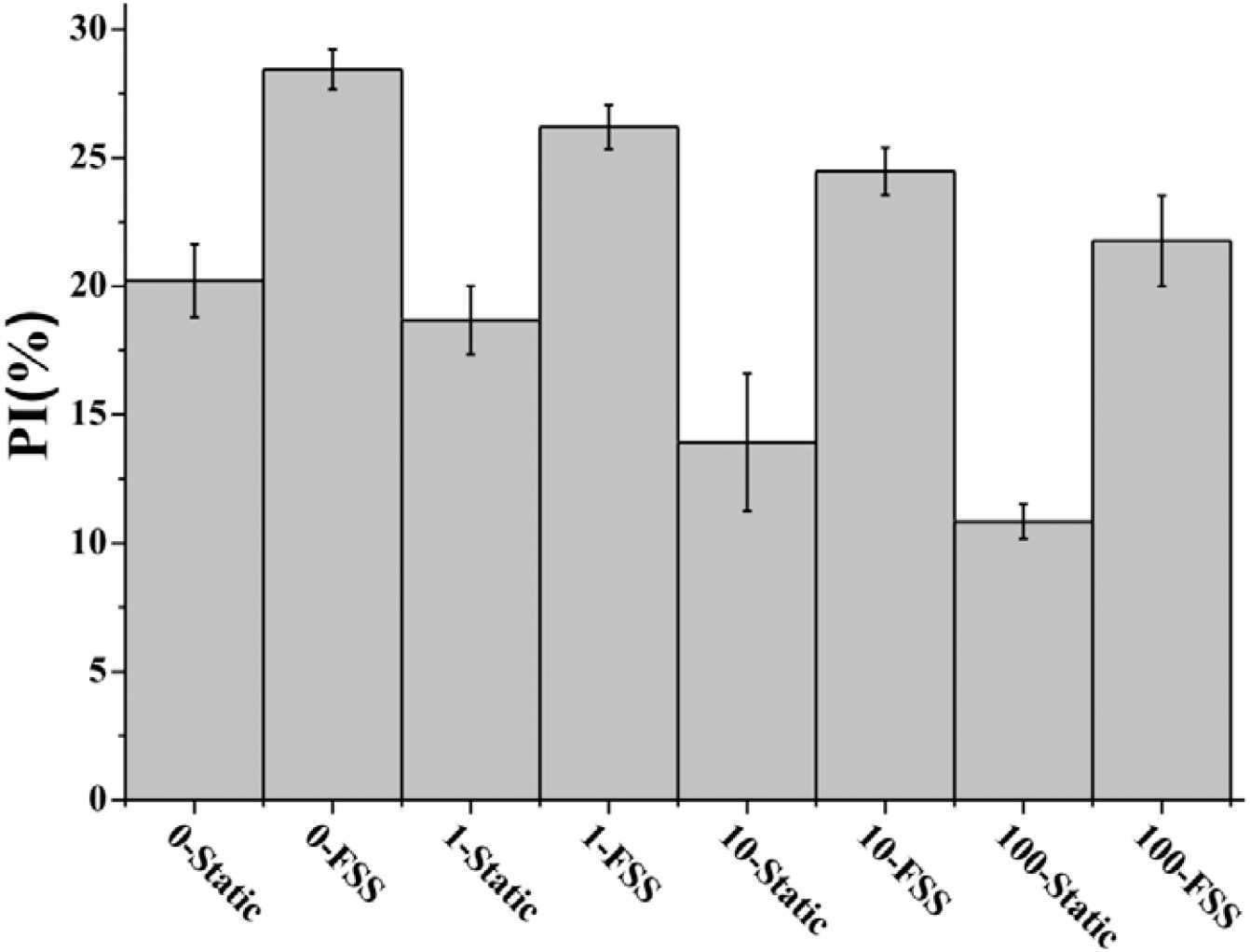
PI values of ROBs with different 1,25 (OH)_2_ D_3_ concentration for Static and FSS.

**Fig. 5.**
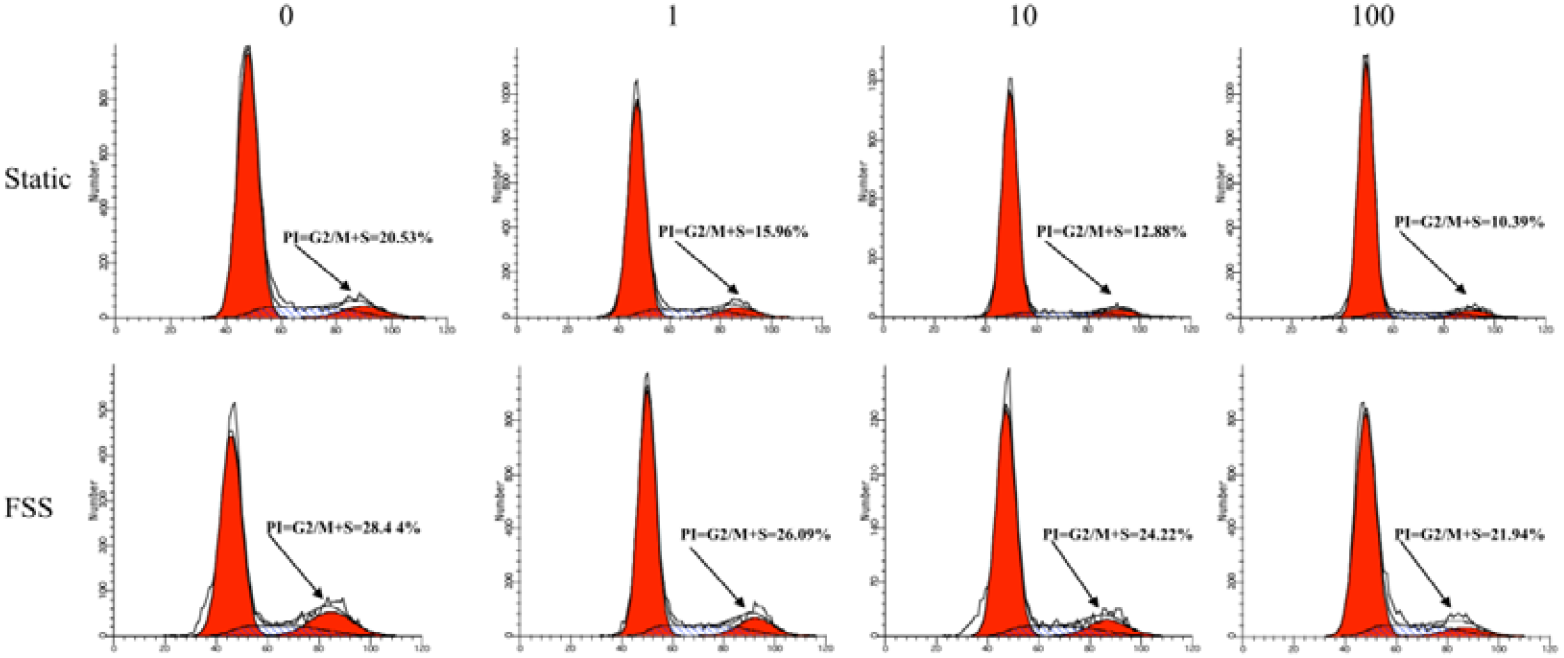
Representative histograms of the PI cell-cycle profiles.

### 3.3. Plasma membrane integrity after FSS exposure

To make sure whether the detected NO and PGE_2_ are actively released by ROBs in response to FSS or passively released due to plasma membrane rupture by FSS, the plasma membrane integrity was monitored by detecting the LDH level in the conditioned media. The ratio of LDH level with FSS exposure to that without FSS exposure was employed to evaluate the plasma membrane integrity. As can be seen from Fig. 6, the LDH ratios for all of FSS-induced ROBs are close to 1. This indicates that the plasma membranes are intact and all NO and PGE_2_ for X-FSS (X=0, 1, 10, 100) are actively released by ROBs.

**Fig. 6.**
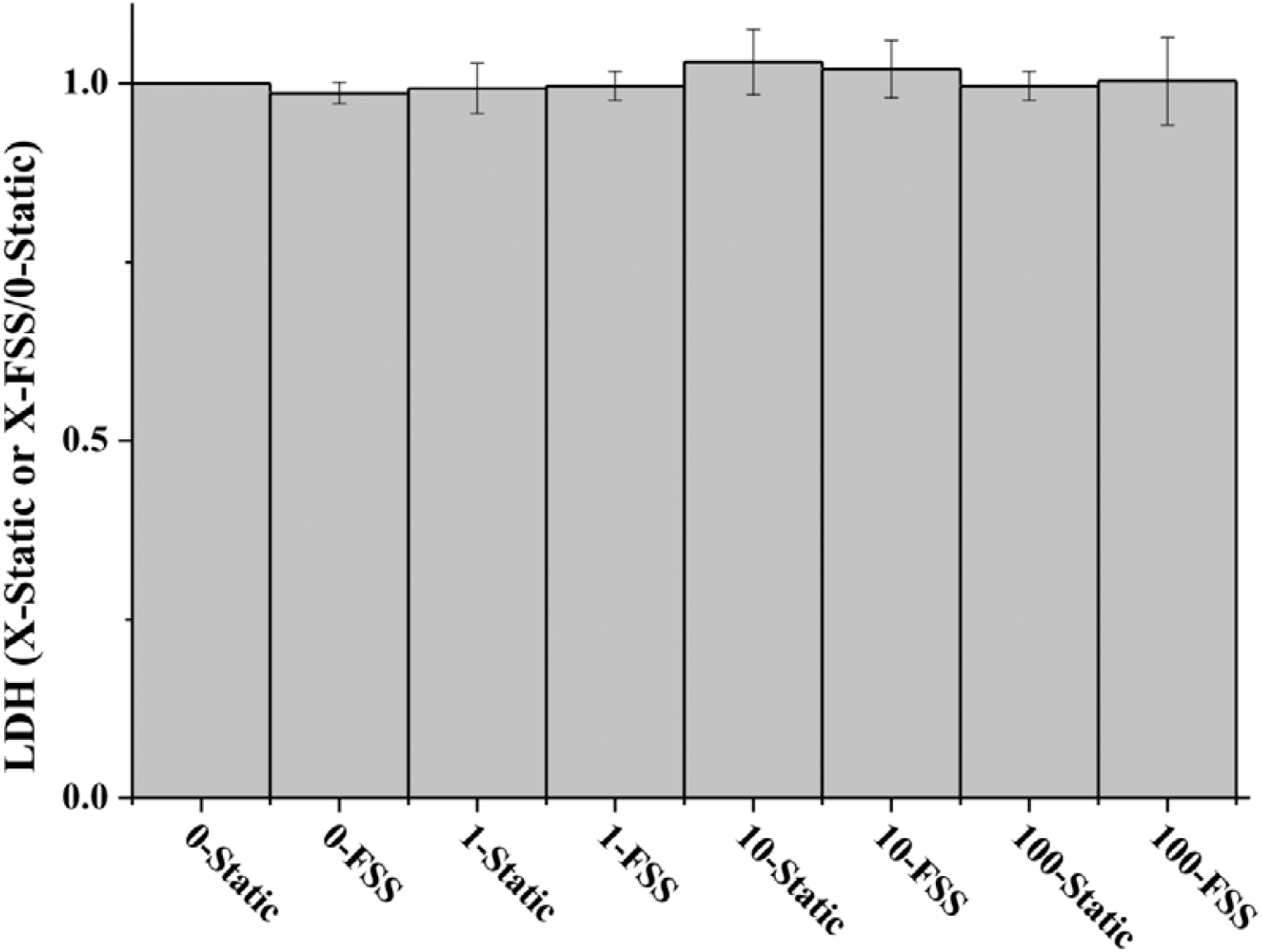
The ratio of LDH release induced for Static and FSS.

### 3.4. Focal adhesions, cytoskeletons and morphologies of ROBs before and after FSS

To elucidate the possible mechanism by which different concentration induces different responses of ROBs to FSS, the focal adhesion formation, F-actin organization and morphologies of ROBs before and after FSS exposure were examined and pictured using a CLSM. The typical pictures were shown in Fig. 7, where the blue, green, and red represent nuclei, vinculin, and F-actin, respectively. Based on the fluorescence images, the cell aspect ratios and angles between cell axis and FSS direction were further measured by using Image J software to quantify cell shape and orientation (Table. 2 and 3). ROBs with an angle of 0-30° are referred to “oriented along FSS direction” while those with an aspect ratio of 1-3 and > 3 are referred to “spread” and “elongated” shape, respectively.

**Fig. 7.**
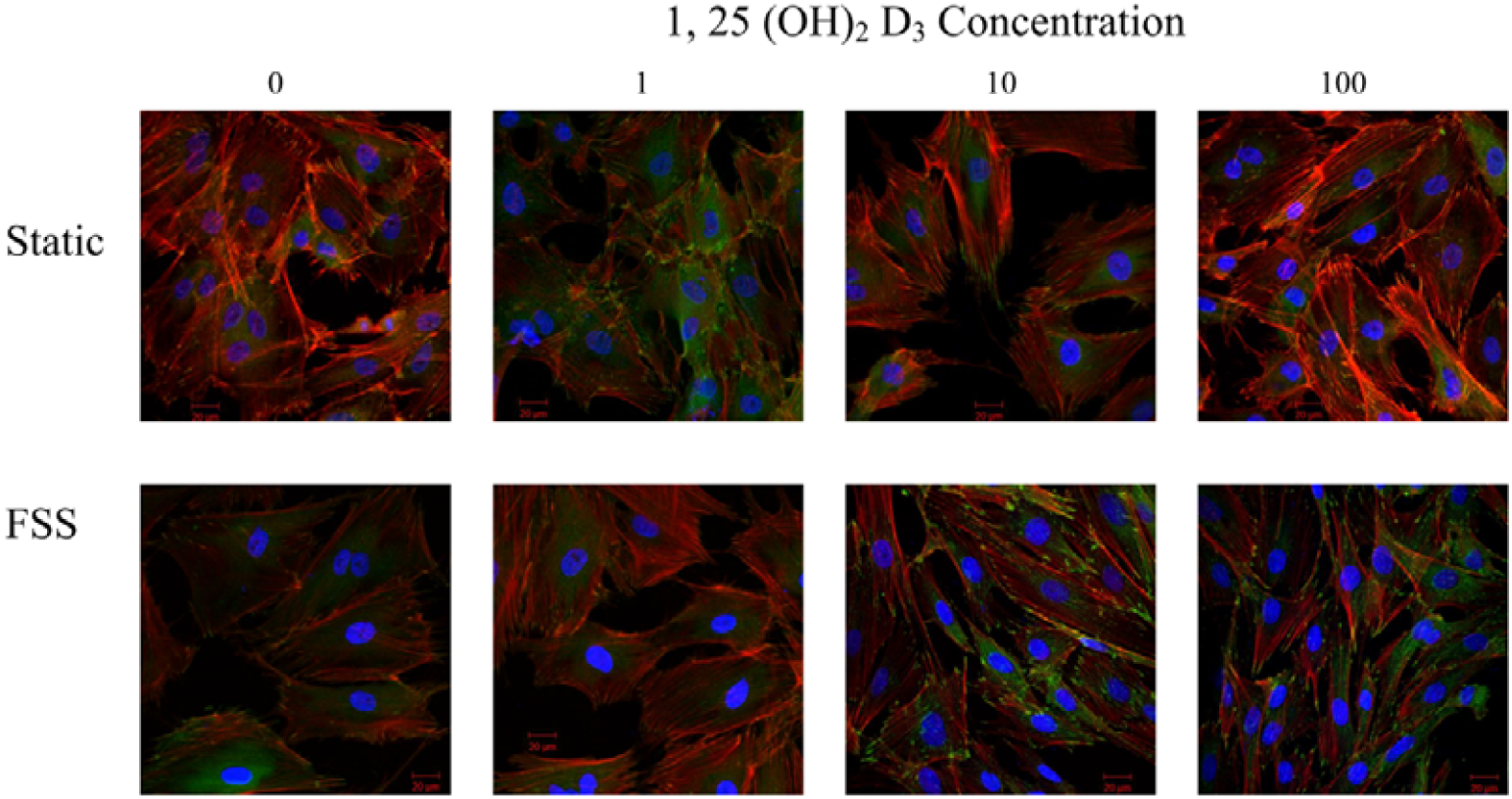
Focal adhesion and F-actin formation of ROBs with different 1, 25 (OH)_2_ D_3_ concentration for Static and FSS. ROBs were fixed, permeabilized, and double labeled for vinculin (green), F-actin (red), and nuclei (blue). The yellow areas are vinculin, partially colocalized with F-actin in the focal adhesion plaques.

**Table 2.**
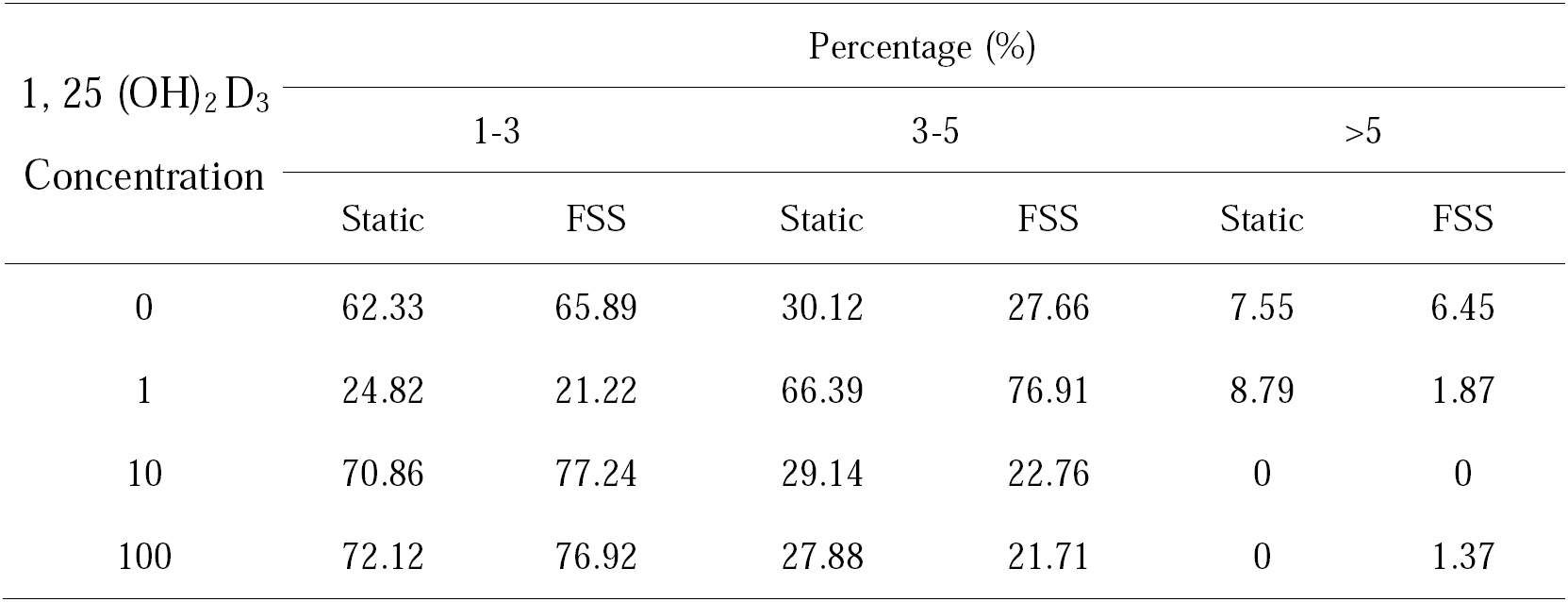
Cell aspect ratios with different 1,25 (OH)_2_ D_3_ concentration for Static and FSS

| 1, 25 (OH)<sub>2</sub> D<sub>3</sub><br>Concentration | Percentage (%) |  |  |  |  |  |
| --- | --- | --- | --- | --- | --- | --- |
|  | 1-3 |  | 3-5 |  | >5 |  |
|  | Static | FSS | Static | FSS | Static | FSS |
| 0 | 62.33 | 65.89 | 30.12 | 27.66 | 7.55 | 6.45 |
| 1 | 24.82 | 21.22 | 66.39 | 76.91 | 8.79 | 1.87 |
| 10 | 70.86 | 77.24 | 29.14 | 22.76 | 0 | 0 |
| 100 | 72.12 | 76.92 | 27.88 | 21.71 | 0 | 1.37 |

**Table 3.** Cell orientation with different 1, 25 (OH)_2_ D_3_ concentration for Static and FSS

| 1, 25 (OH)<sub>2</sub> D<sub>3</sub><br>Concentration | Percentage (%) |  |  |  |  |  |
| --- | --- | --- | --- | --- | --- | --- |
|  | 0°-30° |  | 30°-60° |  | 60°-90° |  |
|  | Static | FSS | Static | FSS | Static | FSS |
| 0 | 32.42 | 75.18 | 29.44 | 20.92 | 38.14 | 3.90 |
| 1 | 34.76 | 71.34 | 31.58 | 22.63 | 33.66 | 6.03 |
| 10 | 35.69 | 77.42 | 29.46 | 19.85 | 34.85 | 2.73 |
| 100 | 32.31 | 74.87 | 37.15 | 23.69 | 30.54 | 1.44 |

Before FSS exposure, 10-Static, and 100-Static spread batter and contained a larger number of focal adhesions and more clear F-actin stress fibers than 0-Static; nevertheless, 1-Static displayed few focal adhesions and vague F-actin stress fibers (Fig. 7). Judging from the cell aspect ratio (Table. 2), 0-Static, 10-Static and 100-Static were spread (62.33%, 70.86% and 72.12% of ROBs with a cell aspect ratio of 1-3) whereas 1-Static were elongated (75.18% of ROBs with a cell aspect ratio of > 3). Regarding cell orientation, ROBs percentages taking an angle of 0°-30°, 30°-60° and 60°-90° are comparable on all of samples, suggesting random orientation of ROBs (Table. 3).

After 1 h of FSS exposure, the percentage of ROBs taking an angle of 0°-30° dramatically increased for all samples (for 0 from 32.42% to 75.18%, for 1 from 34.76% to 71.34%, for 10 from 35.69% to 77.42% and for 100 from 32.31% to 74.87%) (Table. 3). ROBs present a partially oriented morphology along the FSS direction. At the same time, all samples despite a slight decrease in ROBs percentage taking an aspect ratio of 1-3 except 1 (for 0 from 62.33% to 65.89%, for 10 from 70.86% to 77.24% and for 100 from 72.12% to 76.92%) (Table. 2). The cell aspect ratios and angles suggest that FSS induced obvious cell reorientation and slight shape alteration.

### 3.5. ALP analysis

ALP will significantly increase during the post proliferative phase of osteoblasts, and it’s an early marker for differentiation^[29]^ The ALP activity of ROBs by treated with various concentration 1, 25 (OH)_2_ D_3_ and (with or without) FSS for 1 h are shown in Fig. 8, respectively. As we can see that 10-Static and 100-Static enhanced ALP activity of ROBs compared to 0-Static, and 100-Static > 10-Static, which suggests that 10nmol/L and 100nmol/L 1, 25 (OH)_2_ D_3_ promote ROBs differentiation. On the contrary, ALP activity of 1-Static is slightly decreased, and lower than those of 0-Static, which indicates that 1nmol/L 1, 25 (OH)_2_ D_3_ has a negligible inhibition to ROBs differentiation. After ROBs exposed to FSS for 1h, the ALP activity of 1-FSS is increased, and higher than those of 0-Static, which indicates that the exposure of FSS counteracts the inhibitory effect of 1, 25 (OH)_2_ D_3_ on ROBs differentiation. What’s more, ALP activity of 10-FSS and 100-FSS has a significant increase in contrast to 10-Static and 100-Static. The ALP activity of ROBs exposed to various concentrations of 1, 25 (OH)_2_ D_3_ and FSS followed the pattern 100-FSS > 10-FSS > 0-FSS > 1-FSS. It is in good agreement with the suppressed cell proliferation (Fig. 4) which generally increases cell differentiation^[25, 30]^.

**Fig. 8.**
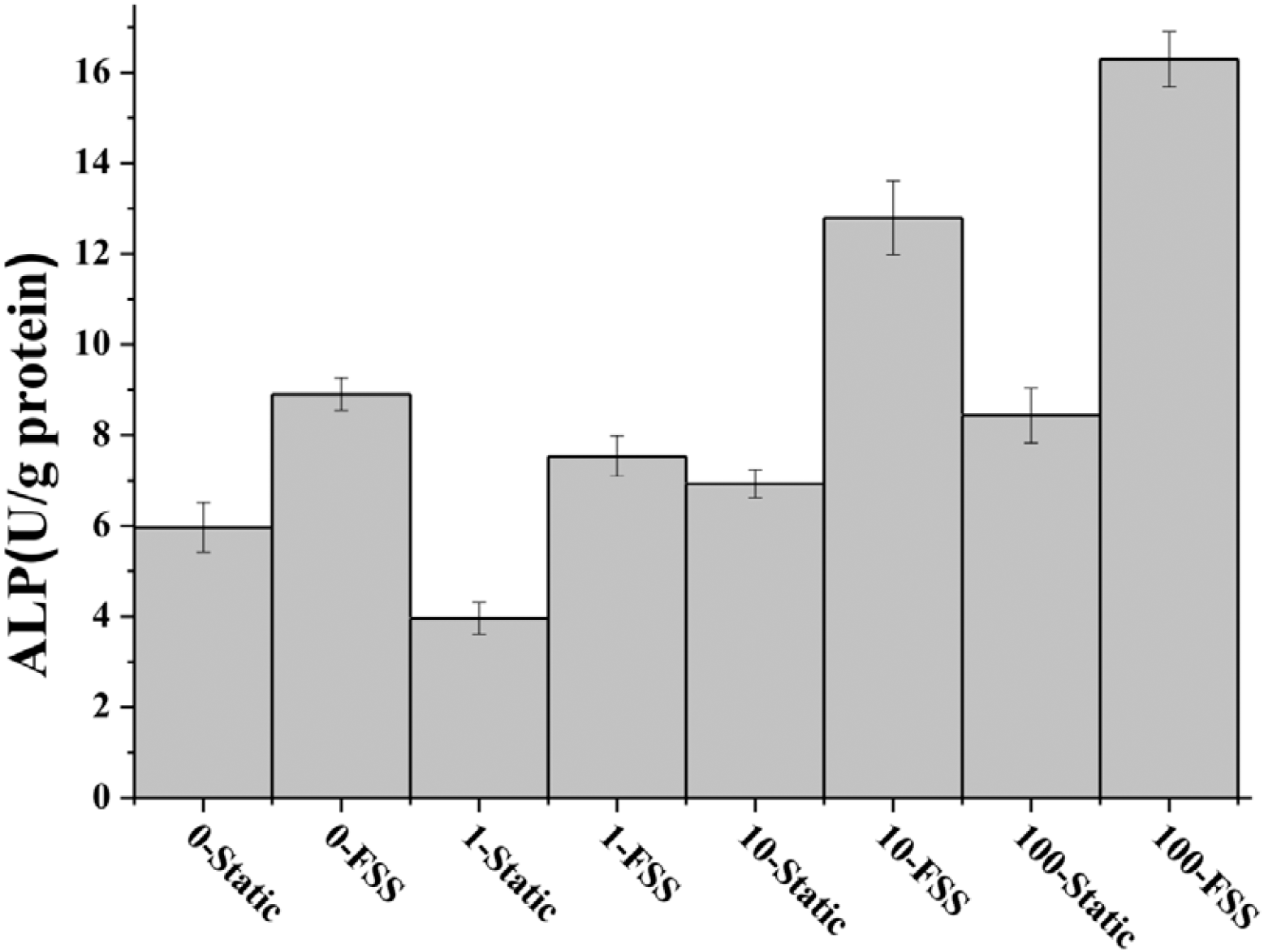
ALP activity of ROBs with different 1, 25 (OH)_2_ D_3_ concentration for Static and FSS.

### 3.6. Quantitative real-time polymerase chain reaction (qRT-PCR)

Fig. 9 summarizes the differentiation-related gene expressions of OPN, OCN for all samples. It is similar to ALP analysis that 10-Static and 100-Static enhanced OPN and OCN gene expressions of ROBs compared to 0-Static, and 100-Static > 10-Static. However, OPN and OCN gene expressions of 1-Static are the same as 0-Static without variation. After ROBs exposed to FSS for 1h, OPN and OCN of 1-FSS is increased, and higher than those of 0-Static, which indicates that FSS enhanced OPN and OCN gene expressions of ROBs by 1nmol/L 1, 25 (OH)_2_ D_3_ treatment. Moreover, OPN and OCN gene expressions of 10-FSS and 100-FSS has a significant increase in contrast to 10-Static and 100-Static. The OPN and OCN gene expressions of ROBs exposed to various concentrations of 1, 25 (OH)_2_ D_3_ and FSS followed the pattern 100-FSS > 10-FSS > 1-FSS ≈ 0-FSS (Fig. 9).

**Fig. 9.**
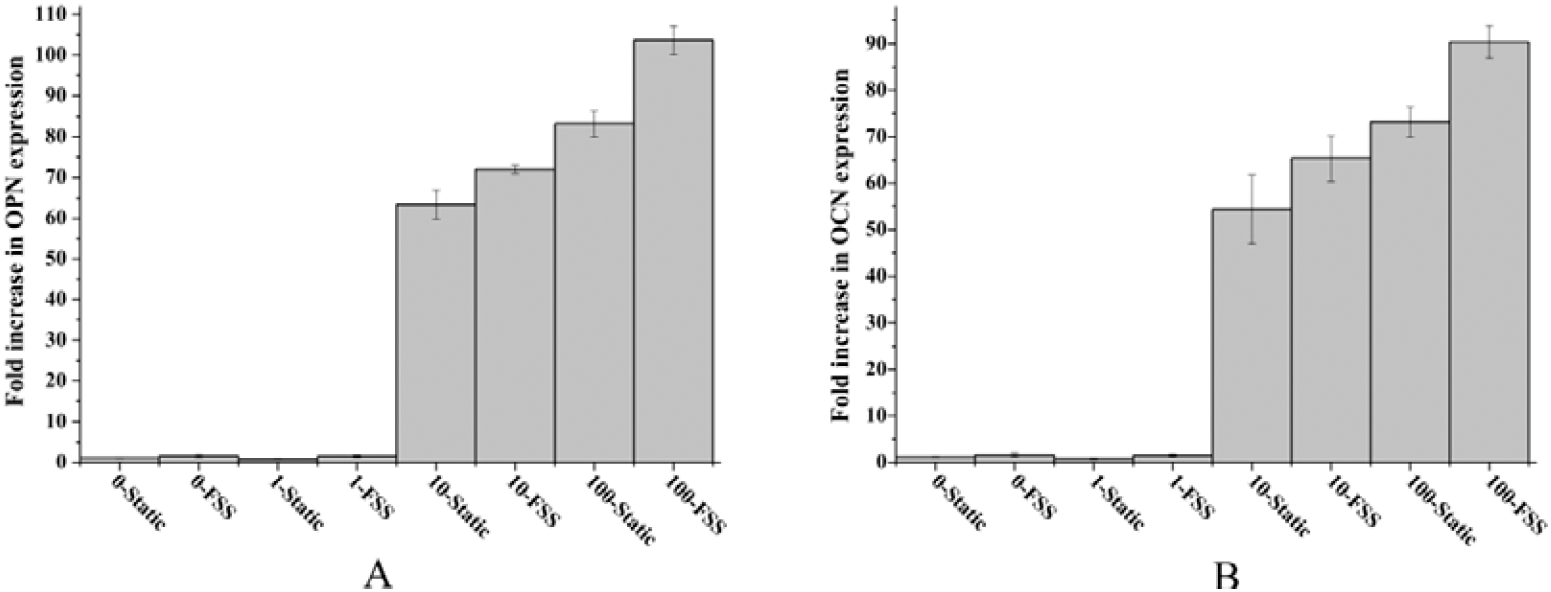
OPN and OCN expression of ROBs with different 1, 25 (OH)_2_ D_3_ concentration for Static and FSS.

## 4. Discussion

The aim of this study was to find the collective effects of 1, 25 (OH)_2_ D_3_ with physiological fluid shear stress on osteoblasts, and to understand the possible mechanism of these responses. 1, 25-dihydroxyvitamin D_3_ and mechanical stimuli are extremely important components for bone tissue engineering. FSS is the principle mechanical stimulus responsible for physiological bone modeling/remodeling^[8–10]^ and thus often employed as an external mechanical stimulus in researching osteobalsts^[31–33]^. In this study, we found that the different concentration of 1, 25 (OH)_2_ D_3_ regulates the responses of osteoblasts to FSS, with the better responses on 10-FSS and 100-FSS yet the worse responses on 1-FSS, than 0-FSS. The potential mechanism is that 1, 25 (OH)_2_ D_3_ modulates the focal adhesions and cytoskeletons which further leads to differential alteration in cell short term responses such as signal molecular (NO and PGE_2_) release, shape, orientation, and long term responses of ALP, PI, and gene expressions such as OPN and OCN after exposed to FSS.

The releases of NO and PGE_2_ are known to play an important role in bone formation and remodeling, and they are essential early responses of osteoblasts to FSS^[34–36]^. In this light, NO and PGE_2_ releases were selected as parameters to indicate the early responses of ROBs. It is not consistent with previous studies about Alone-FSS-induced release of NO^[37–40]^, the collective effects of 1, 25 (OH)_2_ D_3_ with FSS inhibited NO release of ROBs (Fig. 1 and Fig. 2). However, the inhibition of pre-incubation with 1, 25 (OH)_2_ D_3_ to FSS-induced NO release was reported and in line with some previous literatures^[40, 41]^. On the contrary, pre-incubation with 1, 25 (OH)_2_ D_3_ promoted obviously PGE_2_ release of ROBs to FSS, which shown a concentration dependence of 1, 25 (OH)_2_ D_3_ with a pattern of 100-FSS > 10-FSS > 0-FSS > 1-FSS (Fig. 3). Therefore, one conclusion could be made that 1, 25 (OH)_2_ D_3_ inhibited FSS-induced NO release of ROBs, and this inhibition was concentration dependent, the stronger concentration of 1, 25 (OH)_2_ D_3_, the stronger inhibition. At the same time, the high concentration (10nmol/L and 100nmol/L) can promote FSS-induced PGE_2_ release, and low concentration (1nmol/L) reduced.

Furthermore, cell proliferation and differentiation as a long-term response of ROBs was further investigated to evaluate the synergistic effect of 1, 25 (OH)_2_ D_3_ and FSS by using PI, ALP activity, gene expressions of OPN and OCN. We found the same phenomenon that 1, 25 (OH)_2_ D_3_ inhibits osteoblast proliferation consistent with the report of K. Van der Meijden^[5]^. More importantly, the inhibitory effect was concentration dependent like 0-Static > 1-Static> 10-Static> 100-Static (Fig. 4 and Fig. 5). In addition, the addition of FSS eliminated a part of this inhibition, demonstrating that 1-FSS > 0-Static, 10-FSS > 0-Static, 100-FSS > 0-Static (Fig. 4 and Fig. 5). From the perspective of cell differentiation, 1nmol/L 1, 25 (OH)_2_ D_3_ has little effect on ALP activity, OPN and OCN gene expressions, but the addition of FSS appear a remarkable increase for these parameters. This phenomenon illustrates that low concentration of 1, 25 (OH)_2_ D_3_, such as 1nmol/L, could not promote cell differentiation. However, 1-FSS has a remarkable increase on cell differentiation, which shows that FSS plays a dominant role on the synthesis of 1nmol/L 1, 25 (OH)_2_ D_3_ and FSS in cell differentiation. Additionally, the high concentration of 1, 25 (OH)_2_ D_3_ (1nmol/L and 10nmol/L) can promote cell differentiation, and this effect is further enhanced by the intervention of FSS (Fig. 8 and Fig. 9).

Initial adhesion and spreading of osteoblasts, including integrin-medicated cytoskeleton rearrangement^[20–22, 42, 43]^ and initial formation of FA^[44–47]^, have been regarded as critical factors that influence FSS-related mechanotransduction. Pavalko *et al*. proved that FSS-induced mechanical signaling in MC3T3-E1 osteoblasts requires cytoskeleton-integrin interactions^[48]^. However, McGarry *et al*. disrupted the cytoskeleton of osteoblasts and found that FSS-induced NO releases need intact actins, whereas PGE_2_ releases are independent on actin cytoskeleton reorganization^[49]^. In addition, Ponik and Pavalko used bovine serum albumin and RGDS to inhibit formation of focal adhesions in different degrees and confirmed that the releases of FSS-induced osteoblast PGE_2_ are controlled by focal adhesion^[50]^. Taking all above information in mind, one hypothesis could be made that FA and F-actin organization should be responsible for the collective influence of 1, 25-dihydroxyvitamin D_3_ with physiological fluid shear stress on osteoblasts.

To verify this hypothesis, FA and F-actin before FSS exposure were pictured (Fig. 7). The trend of F-actin is 100-Static > 10-Static > 1-Static ≈ 0-Static, while that of the FAs is 100-Static > 10-Static > 1-Static ≈ 0-Static. The common influences of FSS and 1, 25 (OH)_2_ D_3_ revealed that the PGE_2_ releases, ROBs proliferation, ALP activity and gene expressions of OPN and OCN followed the similar patterns of FAs or F-actin organization. These results imply that 1, 25 (OH)_2_ D_3_ regulates the responses of ROBs to FSS by controlling the formation of FA and the organization of F-actin. 100nmol/L1, 25 (OH)_2_ D_3_ produced the best FAs and F-actin organization, so that the optimal combination of 1, 25 (OH)_2_ D_3_ with 12 dynes/cm^2^ physiological FSS was observed on 100nmol/L1, 25 (OH)_2_ D_3_.

In summary, 1, 25 (OH)_2_ D_3_ affects the response of ROBs to FSS, including the inhibition of NO releases and cell proliferation as well as the promotion of PGE_2_ releases and cell differentiation. These findings provide a possible mechanism by which 1, 25 (OH)_2_ D_3_ influences osteoblasts responses to FSS and may provide guidance for the selection of 1, 25 (OH)_2_ D_3_ concentration and mechanical loading in order to *in vitro* produce functional bone tissues.

## Acknowledgements

This work was supported by Grants from the Science and technology support program of Taizhou (No. TS301637), and Innovative Research Team of Taizhou polytechnic college (No. TZYTD-16-4).

